# Developmental Transcriptional Model Describing Regulated Genes, Qtls And Pathways During The Primary And Secondary Cell Walls Of Pima Fibers

**DOI:** 10.1101/056127

**Authors:** Magdy S. Alabady, Bulak A. Arpat

## Abstract

*Gossypium barbadense L*. (Egyptian and Pima) produces single celled fiber trichomes that are the longest and richest in cellulosic contents in the plant kingdom. Developmental dissection of fiber at the transcriptional level is crucial to unveiling the genetic mechanisms underpinning fiber morphogenesis. We profiled the transcriptome of developing Pima fibers, as well as genes associated with consensus fiber quality QTLs, at seven developmental time points covering both primary (PCW) and secondary (SCW) cell wall stages. A total of 2,934 genes were differentially expressed at only one (45.19%) or at multiple (54.81%) developmental time points. Based on the coincidence between gene expression dynamics and the time frame of fiber developmental stages, five stage-specific expression profiles were identified. As a link between fiber QTLs and gene expression, 5 potential developmentally regulated QTLs (drQTLs) corresponding to different fiber developmental stages were identified. Genes in the ubiquitin proteolytic pathway, particularly QTL associated genes, appeared to be involved in regulating the transition stage between PCW and SCW; a stage that is crucial to both fiber length and strength in the extra-long staple cotton genotypes. In this respect, Yeast-two-hybrids identified interactions between UBC9 and genes involved in cell and organ elongation, polar cell expansion, microtubule cytoskeleton dynamics and organization, and basic amino acids transportation during the SCW/SCW transition. Altogether, these results were integrated into a proposed model linking fiber developmental stages with the Pima fiber traits.

## Introduction

Cotton fiber is an epidermal ovule trichome with several distinguishing characteristics making it a unique structure in the plant kingdom. It is the longest single cell and the richest in cellulose content. In addition, cells on the same ovule are synchronously grown and do not undergo cellular division. In the genus Gossypium, there are two major allotetraploid species: *Gossypium hirsutum* L. and *Gossypium barbadense* L. (2n=4x=52, AD genome). In contrast to the high-yielding *hirsutum* cotton (Upland), *barbadense* cotton (Egyptian and Pima) has vastly superior fiber qualities such as length, strength, and fineness, and is highly valued, extra-long staple cotton. The fiber transcriptome of the diploid species *G. arboreum* L., a fiber-producing ancestor of the tetraploid species, was estimated at ~18,000 genes, leading to an approximate prediction of 36,000 homoeologous loci from AD-genome in tetraploids [1]. Approximately, 75-94% of the total cotton genome is transcribed at each of the fiber developmental stages [2]. Fiber genes account for as much as 45-50% of the whole cotton genome fiber genes function is highly conserved, despite millions of years of evolutionary history [3–5]. At the stage-specific level, 12% and 14% of fiber genes are highly active during primary cell wall (PCW) and secondary cell wall (SCW) synthesis, respectively, in developing Upland fiber [1, 6]. The development of terminally differentiated fiber cells occurs in four overlapped stages: differentiation, polar elongation/PCW expansion, SCW synthesis, and maturity [7, 8]. The commencement and duration of these stages are genotype-dependent and influenced by the environmental conditions. The differentiation stage occurs around the time of anthesis (−3 to 1 days post anthesis (dpa)) and is characterized by enlargement and protrusion of ovule epidermal cells that are destined to be fibers [9]. The differentiated fiber cells undergo a vigorous unidirectional elongation and PCW expansion driven by the cell turgor pressure [10] at ~5dpa until reaching the final length at approximately 25dpa [11, 12]. The primary cell walls of cotton fibers are of roughly the same composition as a typical dicot primary cell wall [13]. During SCW construction (~20-45 dpa), the cellulose biosynthesis predominates until the cellulose constitutes more than ~94% of the fiber SCW [11]. The last stage in cotton fiber development is maturation, which continues up to 50 dpa [11]. The biogenesis of cotton fiber PCW and SCW directly affect fiber quality, polygenic traits such as length, strength, fineness, and maturity, where fiber length and fineness are controlled by the genetic activities driving the rate and duration of PCW expansion, while both the rate and duration of SCW synthesis and deposition control fiber traits such as fineness, strength and maturity. In the past two decades, quantitative trait loci (QTL) have been used in conjunction with linkage mapping to identify loci that are primarily responsible for variation in the phenotype of complex, quantitative traits [14]. A major limitation of QTL data is that different parental combinations and/or experiments conducted in different environments often result in the identification of partly or wholly non-overlapping sets of QTL [15]. Nevertheless, more than 400 QTLs related to fiber quality have been identified and mapped using different kinds of mapping populations [15]. QTL mapping using the inter-specific populations (*G. hirsutum* × *G. barbadense*) revealed that although *G. barbadense* parent has fewer fiber QTLs than *G. hirsutum* parent, the majority of the favorable alleles came from the *G. barbadense* parent [16, 17] and that most the mapped QTLs, influencing fiber quality and yield, are located on the “D” sub-genome, which may partly account for the superiority of fiber traits in tetraploid to those in diploid species [18]. However, chromatin transmission from the *G. barbadense* donor was largely and widely deficient [16, 17]. This skewed chromatin transmission is best accounted for by multilocus epistatic interactions [16]. As a result, the introgression of Pima alleles via conventional breeding from inter-specific crosses has not proven successful. This led to the adoption of molecular and functional genomics approaches for improving fiber quality.

The ‘genetical genomic’ approach is considered a key to opening the ‘black box’ that lies between genotype and phenotype, especially with the availability of high-throughput genomics tool. In the past few years, genetical genomics [19] has been used to study the genetic basis of gene expression. The importance of understanding the genetic basis of gene expression is predicated on the idea that the genetic contribution to phenotypic diversity originates from variation in both protein abundance and altered functional changes resulting from mutation [20]. Quantitative geneticists have therefore become very interested in studying the heritability of transcription and detection of expression QTL or eQTL [21]. Transcriptome profiling in systems that are similar to fiber cell has been used, including, for example, the cambial meristem tissue in wood plants [22], the Arabidposis pollen cell [23], the quiescent-center cells of developing roots [24], and vascular tissue epidermal cells in maize [25]. In cotton, microarray has been used to profile the transcripome during the initiation stages of Upland fibers [26] and during the primary and the secondary cell wall stages of Upland fibers [2, 6], at five time points in developing sea-island cotton [27]. Additionally, microarray data-mining was employed to identify Pima alleles contributing to the superiority of Pima fibers relative to upland fibers [28].

In this study, we used a functional genomics approach to profile the transcriptome of developing Pima fiber. We identificated of five stage-specific groups of genes, clustering the developmental expression of genes mapped to fiber quality QTLs revealed five developmentally regulated QTLs (drQTLs). Based on these results, we proposed a model describing the genes, metabolic activities, and drQTLs underlying three developmental stages in Pima fiber.

## Results

### Growth Kinetics of Pima Cotton Fiber

The rate and duration of fiber developmental stages are genotype-dependent and subject to environmental influences. It is therefore necessary to determine the growth kinetics of Pima fibers a *priori* to establish a developmental framework for transcriptome profiling. Under the prescribed greenhouse regime, fiber morphogenesis in Pima S-7 requires ~55-60 days to develop, commencing on the day of anthesis (0dpa) (Figure IA) to boll dehiscence (Figure 1B), at which time the fibers desiccate to form flat, twisted ribbons (Figure 1C and D). Fiber elongation and expansion occurs over a prolonged period of 21 to 25 dpa. Shortly after the onset of fiber morphogenesis (“initiation”), developing fibers undergo linear increase in length after 5dpa, reaching a plateau by 21dpa, by which time fibers have attained ~90% of the final length (Figure 1E). The growth rate increased rapidly and reached a maximum rate of elongation equal to 2.3 mm/day at ~ 9.5dpa (Figure 1F). Termination of the fiber elongation phase occurred in two discrete steps (Figure 1E): a sharp decrease in the growth rate from 10 to 14dpa, and a steady decease from 14 to 21dpa. The latter stage corresponds to the transition stage between PCW elongation and SCW deposition.

**Figure 1:**
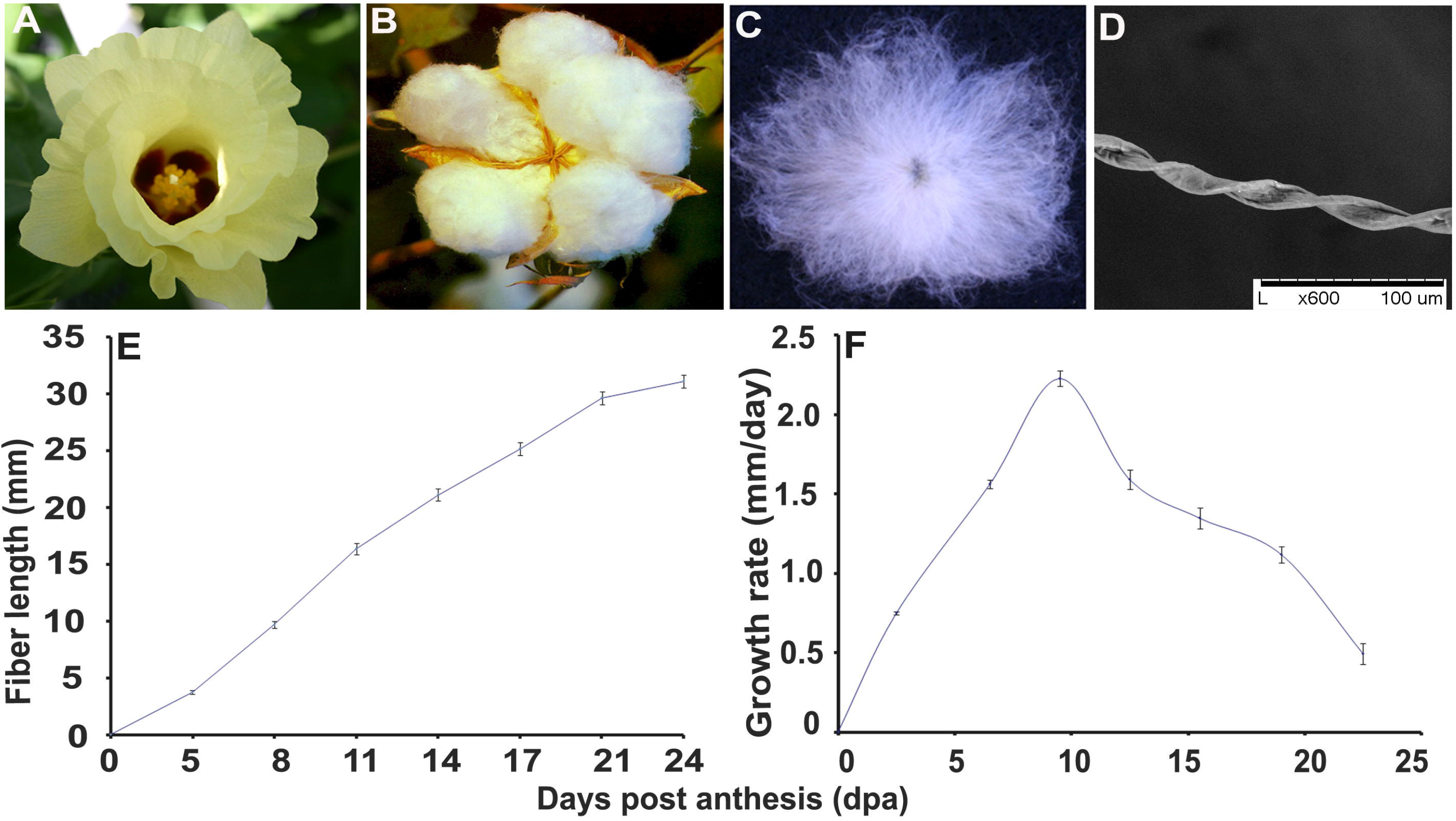
Fiber growth kinetics of extra-long staple cotton (*G. barbadense* L. cv. Pima S7). (A) Pima flower at day of anthesis (o dpa) and commencement of fiber morphogenesis. (B) Mature cotton boll (~55 dpa). (C) Individual seed with >20,000 single-celled lint fibers ~38 mm in length. (D) Fiber growth curve of rapidly elongating Pima fibers. (E) Rate and duration of fiber polar elongation. The peak rate in fiber elongation of 2.3 mm/day occurs at ~9.5 dpa.

### Microarray Data Quality and Validation

The 28 microarray hybridizations produced 1,421,951 data points for 12,227 genes. Detailed description of data processing and quality assessment was published in [28]). Briefly, data quality was improved by the application of four filtration steps, which eliminated 12.7% of the data points, whereas 92.15% (11,267 genes) of the genes with more than 50% of their data points passed all filters. A Strong correlation coefficient between self-control hybridizations was obtained, which demonstrated excellent control of biological variability and dye-bias. In addition, the linearity between direct and indirect hybridizations indicated the reliability of the adopted experimental design and data reproducibility. Quantitative real time PCR (QRT-PCR) was employed to test the accuracy of microarray expression data of 15 randomly selected genes at four time points (8, 14, 17, 21dpa) (Figure S1.) Irrespective of whether genes were lowly or highly expressed, profiles from both techniques were comparable. For up-regulated genes, the Pearson correlation coefficient between data points of each gene from both sources ranged from 0.779 to 0.998. On the other hand, there was a lower correlation coefficient between down regulated genes in microarray and qRT-PCR data. Generally, these results strengthen our confidence in the biological interpretation of the results (Figure S1.)

### Expression profiling of developing Pima fiber transcriptome

We identified 2,934 differentially expressed genes (*P* ≤ 0.05) at one or more time points during the morphogenesis of Pima fiber cell (Table S1). Approximately 45.19% (1326 genes) were differentially regulated in a stage-specific manner, whereas 54.81% (1608 genes) were regulated at more than one-time point, with stages-overlapped (Table 1). With fold change ≥ 2 and FDR-corrected P-value ≤ 0.05, the number of up-and down-regulated genes increased with the progression from PCW to SCW stages (Table 1). We observed a global expression switch at 14dpa in all expression patterns suggesting the commencing point of SCW synthesis, which is accompanied by the declining expansion of PCW and its associated cellular processes (Fig. 2). As illustrated in Figure 2, ten discrete clusters of co-expressed genes relative to the expression at 5dpa were identified. The clusters showing similar expression dynamics but different in the amplitude of the expression ration were joined together to form five stage-specific developmental profiles designated as Type-1: early-expansion associated, Type-2: PCW, Type-3: SCW, Type-4: PCW/SCW transition and Type-5: oscillating.

**Figure 2:**
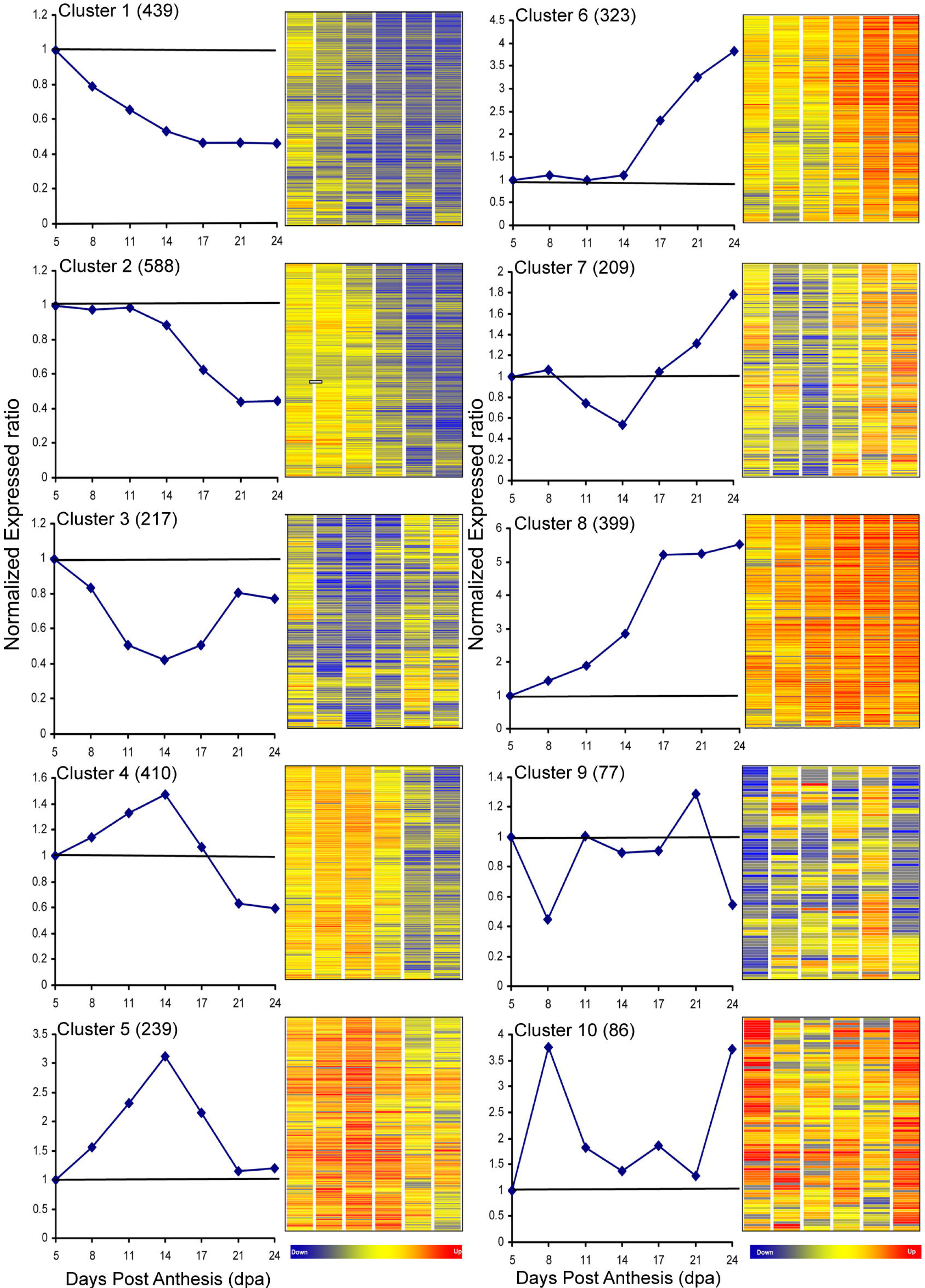
Expression profiles of Pima cotton fiber cell transcriptome during elongation and secondary cell wall synthesis. The k-mean clustering of significantly expressed revealed 10 clusters **(clusters 1 through 10)** with 0.95 cut-off value explaining 90% of the expressional variability within the significantly expressed genes. The expression dynamics of individual genes within each cluster are illustrated by the gene trees. The trend each cluster expression regulation during the PCW and SCW are showed by the graph plot of cluster-average expression.

**Table 1:**
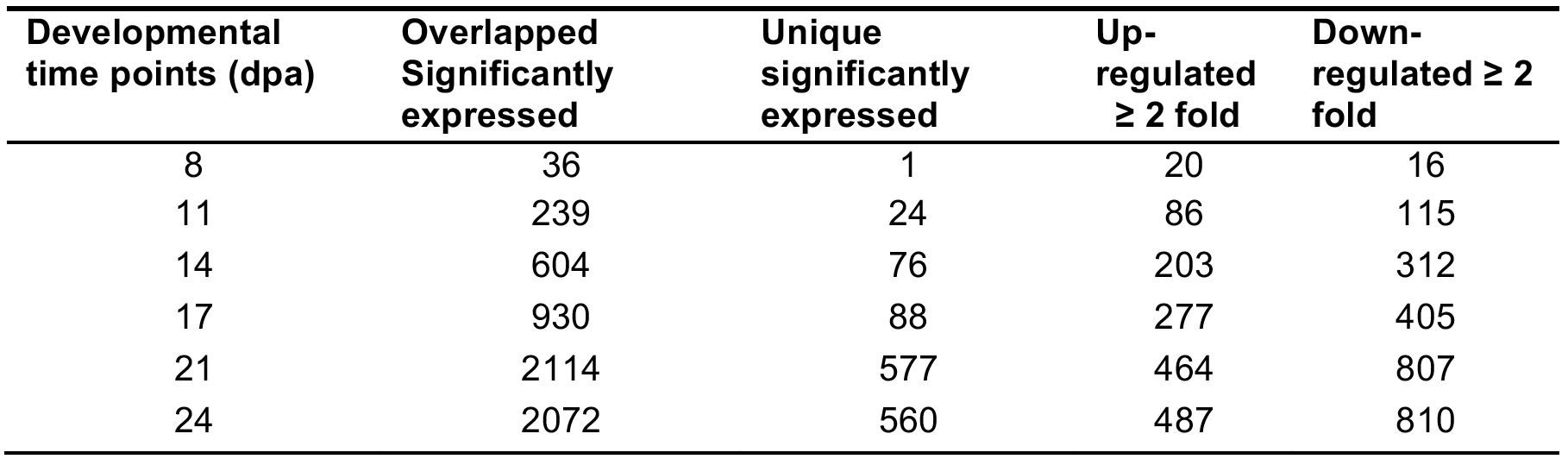
Distribution of the significantly differentially expressed genes (p ≤ 0.05) among six developmental time points in the developing Pima fiber, including genes overlapped and unique as well as up-and down-regulated genes at each time point.

### Type-1: Early-expansion associated genes

During the period from 0-5dpa, fibers enter a phase of early expansion in which cells are slowly expanding and preparing for the commencement of the fast polar elongation stage. Type-1 included all down-regulated genes during both elongation and SCW synthesis stages relative to their expression at 5dpa, and, therefore, was speculated to be associated with the early-expansion phase. Type-1 genes represented 42.40% of the developmentally regulated genes during the morphogenesis of Pima fiber, indicating the complexity of this stage compared to the other developmental stages. Type-1comprised 3 clusters of co-expressed genes named clusters 1, 2, and 3 (Figure 2). Although these clusters were considered as early-expansion specific, they showed different expression dynamics during PCW and SCW stages. In particular while genes in both cluster 1 and 3 were down regulated after 5dpa, only cluster 3 genes were reverted after 14dpa; genes in cluster 2 were down regulated after 11dpa. Among the Type-1 genes were genes (3 in cluster 1 and 2 in cluster 2) active in the phenyl propanoid pathway and ribosomal protein genes (cluster 2) (Table S2). In cluster 3, isoforms of sucrose synthase gene (SuSy), fiber-specific β -1,3-glucanase gene (*GhGlucI*), cellulose synthase-like protein D4 (*Csl*), and 2 putative arabinogalactan-proteins (AGPs) encoding genes were identified. The activities of these genes during fiber initiation and early expansion stages were previously reported [29–33]. The role of cellulose synthase in fiber initiation is still unknown. However, the co-regulation of SuSy, *Csl*, and cellulase genes might suggest a possible role in regulating the plasmodesmata gating in a similar way to the proposed action of *GhGlucI* [31].

### Type-2: PCW specific genes

During fiber elongation, PCW rapidly expand unidirectional by the diffuse growth mechanism [9] until reaching final fiber length. Type-2 included all elongation-associated genes that were up regulated during fiber rapid elongation stage. PCW specific genes were profiled in clusters 4 and 5 (Figure 2) and contained 22.12% (649 genes) of the developmentally regulated gene. While the highest expression levels in this category were reached by genes of unknown function or hypothetical proteins, most of what have previously been described as elongation associated genes were in Type-2 (Table S2). For instance, genes that are known to be vital in maintaining turgor pressure, such as fiber-specific acyl carrier protein, ACP [34], vacuolar H^+^-ATPase subunit D [35, 36], plasma membrane H^+^-ATPase [37], and tonoplast intrinsic protein [38] were 3 fold up regulated at 8 and 11dpa, coinciding with the peak of Pima fiber growth rate. Similarly, four tubulin isotypes were up regulated: β-tubulin Chain-1, Chain-8, Chain-9 and α-tubulin Chain 1. Not only were β-tubulin Chains 8 and 9 the highest up regulated isotypes, but they were also more abundant than α- tubulin isotypes during the elongation stage of Pima fiber. In cotton unigenes, there are eight expansin genes, and they were all represented in the microarray platform used in this study. Only two α- and one β-expansins were up regulated in the elongated Pima fiber. However, the two α-expansins, homologous to *Arabidopsis Exp4* and *Exp3*, were more abundant than the β-expansin, which is homologous to *Arabidopsis Exp2*. Out of eight sucrose synthase members (SuSy) in the known portion of fiber transcriptome, SuSy isoform was the second highest up-regulated gene within PCW specific genes, the peak expression occurred at 14dpa with an average of 8.3 fold during the interval from 5 to 14dpa (Table S2). In general, the expression of PCW marker genes confirmed the association of Type-2 genes with the PCW as major players in the determination of Pima fiber length.

### Type-3: SCW specific genes

During the SCW stage, cellulose synthesis predominates until the cellulose reaches more than 90% of the fiber contents [11]. Differentially regulated genes after the global expression switch at 14dpa were considered to be SCW specific. Type-3 comprised 18.13% (532 genes) of developmentally regulated genes in the developing Pima fiber and represented in clusters 6 and 7 (Figure 2). Whereas co-expressed genes in both clusters were similarly expressed during SCW, their expression patterns were different during PCW. The only up-regulated cellulose synthase subunit (expressed at 15.65 fold) during SCW was highly similar to the *Gossypium hirsutum* cellulose synthase-like catalytic subunit (*CelA3*) (Table S2). In addition to cellulose synthase, endo-1,4-beta-glucanase (EGase family) is also required in cellulose biosynthesis [39]. Two major EGase isoforms were co-expressed with *CesA* during the synthesis of Pima fiber SCW. The expression of these SCW marker genes reinforces the expression dynamics-based association between Type-3 genes and SCW stage.

### Type-4: PCW/SCW transition stages

In the transition stage PCW expansion gradually declines whereas SCW deposition begins. In developing Pima fiber, coinciding with PCW/SCW transition (14-21dpa), 13.60% were up regulated with expression dynamics depicted in Figure 2 cluster 8. The sudden increase in the expression of Type-4 genes at the PCW/SCW transition stage suggested that these genes were associated with the three major processes taking place at this stage: microtubule re-orientation, cessation of the PCW elongation, and commencement of the SCW cellulose synthesis machinery. As a marker for cytoskeleton re-orientation, profilin was up-regulated 9.37 fold during the PCW/SCW transition stage versus 6.84 fold during the elongation stage. During this transition stage, LIM-like protein, a transcription factor in lignin biosynthesis, was highly up regulated (33 fold) (Table S2). Interestingly, 3 *MYB* genes were up expressed during PCW and SCW with a slightly higher expression level in SCW (Table S2). Additionally, four ubiquitin-dependent proteolytic pathway associated genes were up regulated: 26S proteasome AAA-ATPase subunit RPT5a, 2 Ub-conjugating enzymes (E2), and an E3-SKP1 ligase. SKP1 is a part of the E3-SCF complex (SKP1/Cullin/F-box) which is mediating the ubiquitination of transcriptional repressors in response to auxin, promoting auxin signaling [40]. A gene of unknown function had the highest level of gene expression during Pima fiber development. It was expressed 67 fold during the elongation stage and 827 fold during the transition stage and early SCW stages (Table S2). The Pfam domain analysis of this gene revealed an insignificant similarity to a collagen domain. Another two unknown genes were highly expressed: 23 and 15 fold, respectively. This extremely high level of expression points to these genes as major players in fiber development.

### Type-5: Genes with oscillating expression pattern

This category included 5.55% of developmentally regulated genes. In Cluster 9, a group of 77 genes were expressed in an oscillating manner with a down regulation peak at 11dpa and an up regulation peak at 21dpa (Figure 2). Similarly, cluster 10 (86 genes) has two up-regulation peaks at 11dpa and 21dpa (Figure 2). This category was found rich with genes with predicted regulatory functions, including 21 genes encoding binding proteins: 6 DNA, 5 RNA, 4 proteins, and 6 metal binding proteins (Table S2). In the DNA binding group, there were 2 *Myb*-like transcription factors: *Myb*-like protein and *GhMyb*-like. Both members were highly expressed around the expansion peak, whereas *GhMyb*-like was highly expressed at the peak of cellulose synthesis at 24dpa. The highest up-regulated RNA binding protein encoding gene was similar to rice RNA and export factor-binding protein (*REF1-I*). This gene was up regulated around the peaks of expansion and cellulose synthesis at 9.5 and 24dpa, respectively. In the protein binding group, a WD-repeat protein encoding gene was up regulated during both elongation and SCW deposition with peaks at 8 and 24dpa. Another binding protein encoding gene was expressed in the same pattern as the WD-repeat (Table S2). In the ion binding protein, a putative ubiquitin E3 ligase (RING zinc finger protein) gene was up regulated at 8 and 14dpa. Diphenol oxidase (*Dox*) is a copper binding protein, which was up regulated at 8dpa and from 14 to 24dpa. *Dox* is a probable 26S proteasome non-ATPase regulatory subunit 3, prophenoloxidase (www.ihop-net.org). S-adenosylmethionine synthase, a magnesium ion binding protein, was highly expressed during the peak of cellulose synthesis at 24dpa. Similarly, an iron binding protein, lipoic acid synthase was up regulated at the peak of cellulose synthesis and, also, at the elongation peak. Altogether, Type-5 genes appear to be involved in regulating genes associated with both PCW and SCW in the developing Pima fibers.

### Developmental expression analysis of fiber quality QTL

Fiber quality QTL-associated genes were integrated in the developmentally regulated fiber transcriptome to gain deeper insights into the influences of gene regulation on the fiber phenotype. Initially, bioinformatics-based comparative mapping approach was employed to enrich and develop consensus fiber QTL regions. This approach enriched the 81 QTLs derived from primary QTL map [18] by adding 631 and 861 loci from corresponding chromosomes and homoeologous chromosomes in the linkage map [3], respectively. The resulting level of enrichment was approximately 300%, with 1,992 loci after the enrichment compared to 499 loci before the enrichment. In general, FS QTLs were enriched the most, followed by FF, FL, FE, FC, and FLU (Figure 3A). The overlapping between markers derived from the QTL primary map and Linkage map is illustrated by the Venn diagram in Figure 3B. Out of the genes mapped to 1,992 QTL-associated loci, 429 genes were included in microarray platform and among them 144 (33.56%) genes were developmentally regulated. This percentage is significantly higher than the general expected ratio of significantly expressed gene, 23.99% as estimated in this study (Table S3). This finding suggests that QTL-associated genes are preferentially expressed during the various stages of development and therefore can be used to identify potential QTL networks.

**Figure 3:**
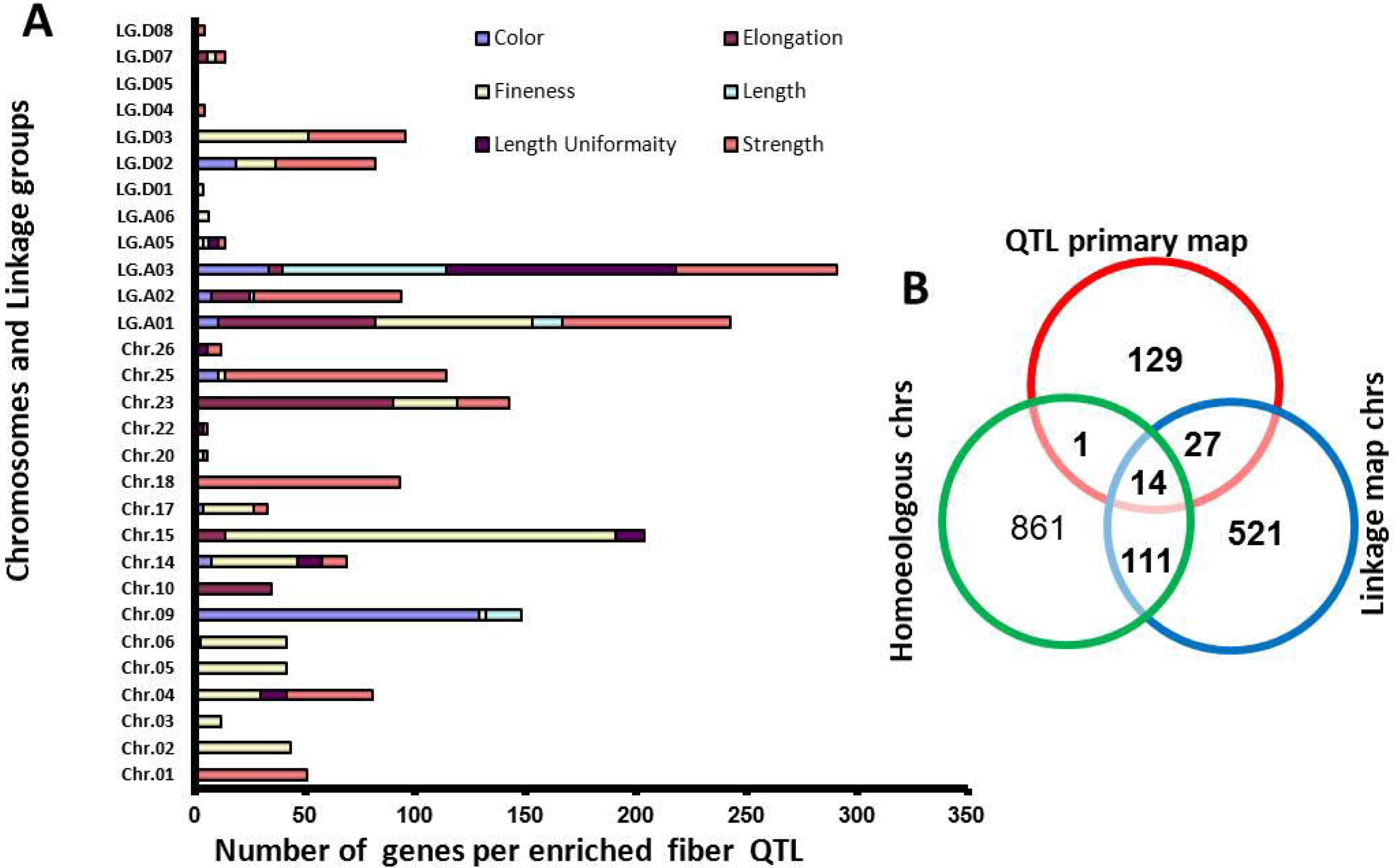
Linking developmentally regulated genes with fiber QTLs. Distribution of enriched fiber QTLs on the cotton chromosomes and linkage groups **(A)** showed superiority of FS and FF QTLs. Enriched fiber QTL by linking to homoeologous chromosomes added 861 gens to the 81 fiber QTLs **(B)**.

### Developmentally regulated fiber QTL networks

Developmentally regulated QTL-associated genes were employed to identify potential QTL networks during different fiber developmental stages. A set of 1,007 genes (50.5%) were overlapped between different QTLs for different fiber traits (inter-QTL overlapping) and among different QTLs for the same trait (intra-QTL overlapping). Another set of 985 (49.5%) markers were uniquely mapped to specific QTLs. A Boolean scoring of the unique and inter-and intra-QTL genes was used to assess the similarity level between QTLs and hence identifying QTL clusters. As illustrated in the UPGMA-based dendrogram (Figure 4A), the lowest similarity level was (68.13) between FC and FS, while the highest was (89.5) betweenFLand FLU. The first subcluster included FL, FLU, FC, and FE QTLs. The other two sub-clusters were assigned for FF and FS QTLs, which were relatively distant from the first sub-cluster. Generally, the inter-QTLs similarity levels were relatively high and ranged from 90% to 68%, which suggests a high level of QTL-QTL interaction during the fiber development. These findings highlight the need for the identification of master QTL networks that have a major effect on fiber development Therefore, potential developmentally regulated QTLs (drQTLs) networks were identified by clustering the expression of QTL-associated genes coinciding with various developmental stages.

**Figure 4:**
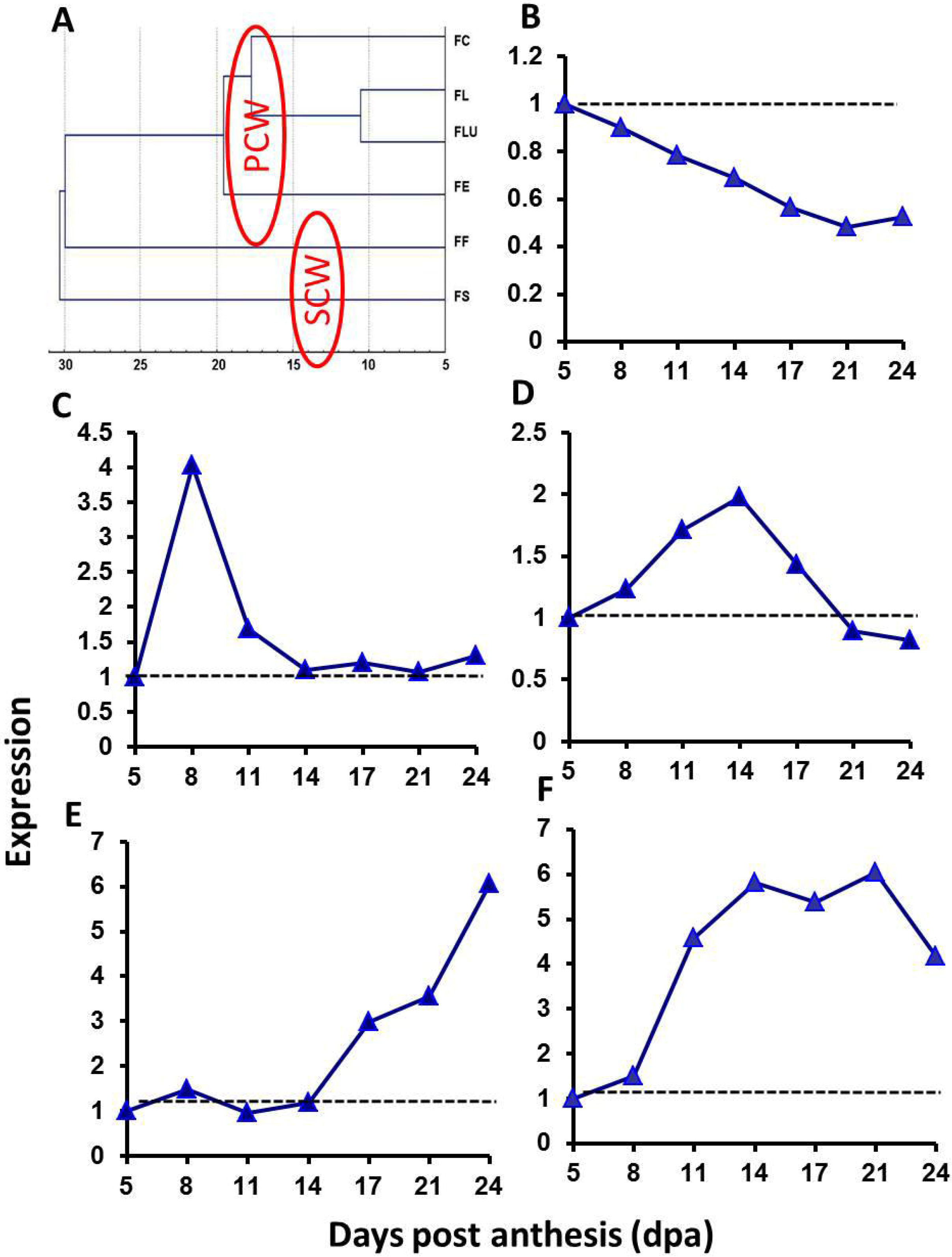
Fiber developmentally regulated QTL (drQTL) networks. **(A)** dendrogram showing the inter-QTLs similarity for six major fiber traits. K-means cluster analysis of the QTL-associated developmentally regulated genes identified 5 potential drQTL networks corresponding to different stages of fiber development **(B** through **F)**. The dotted line represents the expression level at 5 dpa, the reference time point.

### Potential early-expansion drQTL

Although genes in this drQTL were mapped to many QTLs (Table 2), they had synchronized up regulation profiles during the early-expansion stage (Figure 4B). In this drQTL, some of the members were mapped to physically overlapped QTLS on the same chromosome or across homoeologous chromosomes. Specifically, 24 members were mapped to 14 non-overlapped consensus QTLs: 2 FC, 2 FE, 6 FF, and 4 FS QTLs. Whereas 39 members were mapped to 28 overlapped consensus QTLs: 1 FC, 2 FE, 9 FF, 3 FL, 4 FLU, and 9 FS QTLs. The vast majority of the members were originally mapped to QTLs such as FF, FE, and FS. This finding is interesting taking into account that the FF and FS are affiliated more with SCW and that the stages of early-expansion and SCW synthesis are separated by the long elongation stage.

**Table 2:** List of the member genes in the four potential developmental regulated QTL (drQTL) networks

|  | Whole array | QTL | Ubiquitin genes | QTL-Ubiquitin | 26S proteasome |
| --- | --- | --- | --- | --- | --- |
| Microarray probes | 12,227 | 429 | 487 | 30 | 113 |
| Significant | 2934 | 144 | 125 | 15 | 33 |
| %* | 23.99 | 33.56 | 25.66 | 50 | 29.20 |
| ***A. thaliana* matches** | NA | NA | 241 | NA | 50 |

\*% calculated relative to the number of microarray probes of each category

### drQTL active during the peak of fiber elongation

Based on the current saturation level of fiber QTLs, only 2 genes participated in this drQTL (Table 2). Although the expression dynamics of these genes showed peak expression around the fiber elongation peak (Figure 4C), genes were mapped onto two consensus fiber strength QTL: FS25_1 and FS4_1. One of the members is *Myb*-like gene (FS25_1), a transcription factor that has been known to play a major role in fiber trichome initiation and expansion [41]. Obviously, this drQTL contains more genes, which could be discovered upon the saturation of the cotton map or sequencing the cotton genome.

### PCW expansion related drQTL

The expression dynamic of this drQTL was up regulated in coincidence with fiber PCW expansion stage (Figure 4D), which involved elevated activities of members of the expansin gene family. Particularly, β-expansin2 (C0N_001_06916), which was mapped to a position on the LG.A03 overlapping three different QTLs: FLU, FL, and FS (Table 2). Fiber annexin is another marker for the fiber expansion stage and it was included in this drQTL. It was uniquely mapped to the consensus FF14_1 QTL. Annexin plays a role in plant cell development by modulating the activity and/or localization of callose synthase [42]. A set of 32 developmentally regulated genes included in this drQTL. These genes were mapped to 31 consensus fiber QTLs: 3 FC, 3 FE, 11 FF, 2 FL, 2 FLU, and 10 FS; 50% of these genes were overlapped between more than one QTL (Table 2). It is noteworthy that the number of members associated with FF and FS QTLs was higher than the number of those associated with FL QTLs, although it is well-known that fiber length is an expansion-dependent trait while FF and FS are SCW-dependent traits. This notion reinforces the need of dissecting QTLs at the expression level and re-mapping them to saturate the drQTL networks.

### drQTL associated with SCW stage

The expression dynamics of the genes in this drQTL were typical for SCW specific pattern. These genes were expressed slightly above the basal level during the expansion stage followed by an aggressive linear increase in the expression starting at 14dpa, the beginning of PCW/SCW transition stage (Figure 4E). In this drQTLs, 18 genes were mapped to 25 overlapped consensus QTLs: 3 FE, 8 FF, 2 FL, 1 FLU, and 1 FS QTLs (Table 2). Seven genes were uniquely mapped to 5 consensus QTLs: 2, 2, 1, 1, and 1 genes were mapped to FC9_1, FF5_1, FS4_1, FS25_1, and FE10_1 QTLs, respectively (Table 3). A key member in this network, and in SCW deposition process, is membrane-anchored endo-1,4-β-glucanase gene (*CesA*), which was uniquely mapped to the consensus FE10_1 QTL (Table 2). Interestingly, this drQTL included at least 3 Ubiquitin-pathway (Ub-pathway) related genes: 2 UBA-system Cue component genes (FS15_1), and one E3 ligase gene (C3HC4-type RING finger), which was mapped onto two non-overlapped consensus FF QTLs on Chr.15.

**Table 3:** Results of the Yeast-two-hybrids interactions between UBC9 as bait and the developing Pima fiber transcriptome at 114-17dpa

| Bait (pGBKT7) | Prey (pGADT7) | Colony color |
| --- | --- | --- |
| GbUBC9 (#AAR83891) | 26Sproteasome (#AAL32634) | ++ |
|  | GbCHIP (#NP_566305) | +++ |
|  | ATUBA1 (#CAA71762) | ++ |
|  | UBQ4 (#CAJ86369) | +++ |
|  | UBQ10 (#CAA40324) | ++ |
|  | Arginase 1 (#AAV36808) | + |
|  | Acyltransferase (#AAL67994) | +++ |
|  | Dynamin (#ABD28549) | + |
| | β-tubulin5 (#ABE90057) | ++ |
|  | Zinc finger (#ABE89604) | +++ |
| SV40 | Lam | — |
| SV40 | P53 | +++ |

#: GenBank Accession No. Yellow and purple color represents Y2H assay and confirmation by performing Y2H vector swap experiment separately. The intensity of the interaction was measured in the colony color assay. The colony color assay is graded as light blue (+), medium blue (++), or dark blue (+++), (-) represents no interaction.

### PCW/SCW transition drQTL

Members of this network were highly up regulated during the transition stage between PCW and SCW (Figure 4F) and therefore predicted to play major roles in the starting of SCW while ending the PCW stages. In this network, 10 members were mapped overlapping multiple consensus QTLS for different traits whereas 5 members were uniquely mapped to consensus QTLs (Table 3). This network is enriched with genes containing binding domains like WD-40 and Zinc finger, which reflects its regulatory nature. Major feature of this network is the membership of at least 7 Ub-pathway related genes (Table 2). Interestingly, 6 out of the 7 were mapped to fiber fineness (FF) consensus QTLS: FF5_1, FF6_1, FF15_1, and FF17_1. Given that fiber fineness is a function in both fiber length and thickness, this finding is strikingly reinforces our previous speculation that Ub-pathway plays crucial role in regulating the PCW/SCW transition stage.

### Functional categorization and pathway analysis

The developmentally regulated genes of known functions (68.50% of all developmentally regulated genes) were grouped into GO functional categories (*P* < 0.05) and mapped to metabolic pathways. In the structural molecule activities (Figure 5A), 9% of the genes were associated with the structural constituent of the cell wall compared to 1.74% reported in the GO databases (Figure 5 A–1). At the level of catalytic activities (Figure 5 A–2), within the transferase process, the 5-methylthioribose kinase and cellulose synthase were the highest represented gene families. Within the hydrolase processes, cellulase and galactosidase were the top represented families. All gene families belonging to oxidoreductase processes were significantly over represented as well. At the level of binding activities in the fiber transcriptome (Figure 5A–3), the nucleic acid binding process was under represented, while the mRNA and sugar binding processes were highly over represented. The biological processes annotation (Figure 5B) revealed a bias to Ubiquitin-dependent protein catabolism, cellular metabolism, and cell organization and biogenesis. The protein catabolic pathways (Figure 5B-1) in the developing fiber cell were conferred to ubiquitin dependent catabolism in which the proteosomal dependent and ubiquitin modification dependent catabolism were represented by 21% and 79% of the whole catabolic pathway, respectively. Carbohydrate metabolism (50%) and nucleosome assembly (33%) were the top groups in the cellular metabolism and biosynthesis (Figures 5B-2 and 5B-3), respectively. The global functional categories were dissected to identify the major processes at each fiber developmental stage. During the initiation stage, mitochondrion organization and biogenesis was the most over-represented category, whereas mitotic cell cycle category was the most underrepresent (Table S3). At the level of expansion-associated genes, protein biosynthesis and cytoplasm organization and biosynthesis were the highest over-represented functional categories. On the other hand, catabolism activity was the most under-represented functional category during the PCW expansion stage (Table S3). In the cellulose synthesis-associated genes (PCW/SCW and SCW specific), the most over-represented GO category was ubiquitin-dependent protein degradation and carbohydrate transport. Transcription activities were the most under-represented functional category in the SCW specific genes. Common to PCW and SCW the highest over-represented category was G-protein signaling, while the most under-represented category was DNA metabolism. In the oscillating expression gene category, defense response and enzyme regulator activities were the top over-represented functional categories. At the level of pathway analysis, KEGG Ub/26S proteasome proteolytic pathway and ribosome biogenesis and assembly were the top significantly (*P* = 0) regulated pathways. Gap junction (*P* = 1.57e-11) was the third ranked regulated pathway in the developing fiber cell. Flavonoid biosynthesis (*P* = 0.001198), protein folding and associated processing (*P* = 0.0015), nucleotide sugar metabolism (*P* = 0.0051), and mTOR signaling (*P* = 0.0084), a protein kinase involved in regulation of protein synthesis in response to growth factors, nutrients, and energy availability, pathways were developmentally regulated across all developmental stages.

**Figure 5:**
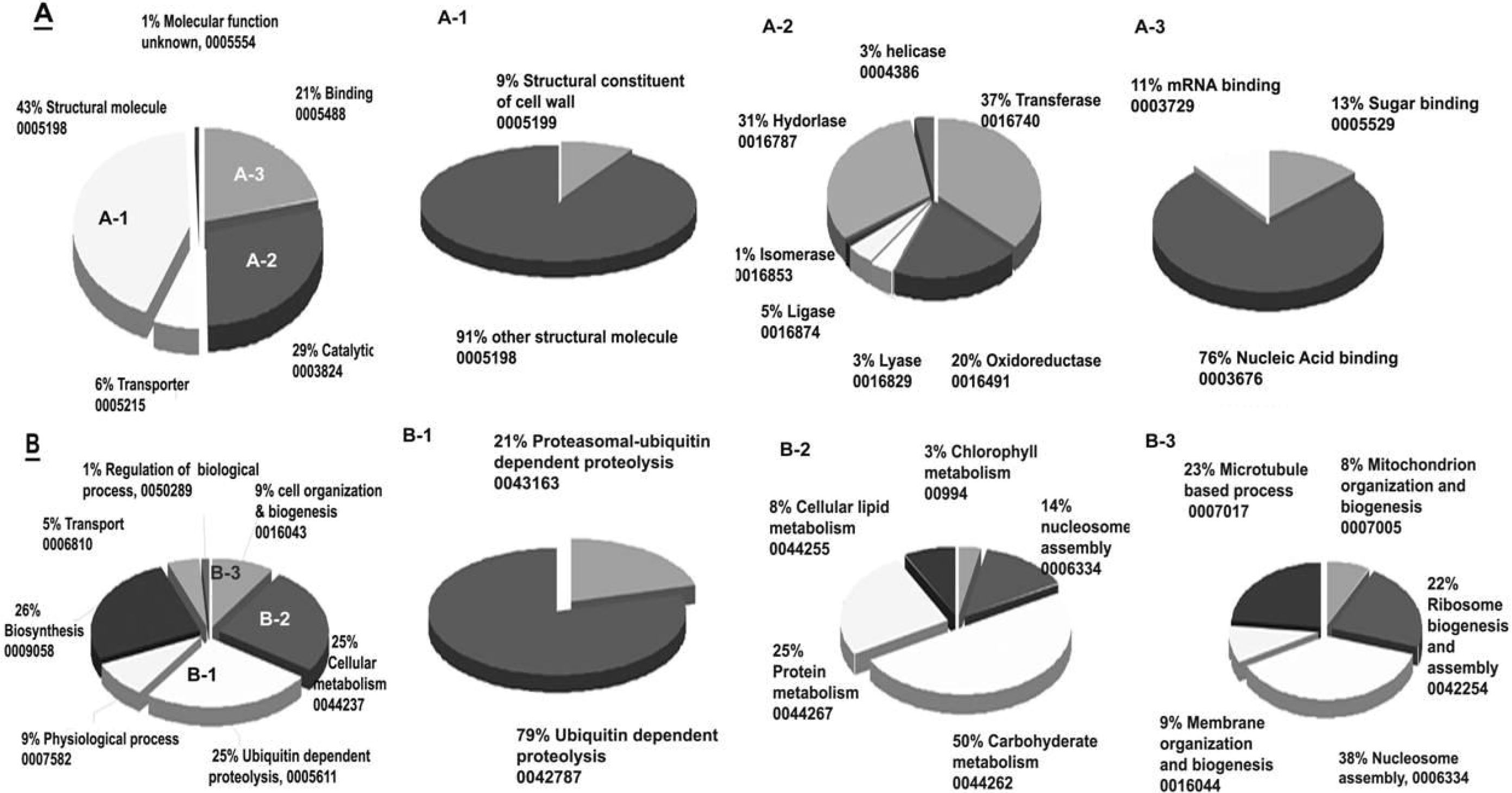
GO categories of significantly expressed and developmentally regulated genes in Pima cotton fiber transcriptome at PCW and SCW synthesis. The percentage of fiber genes that are annotated to Molecular Function and Biological Process GO categories are depicted by the pie charts **A** and **B**, respectively. The Major sub-classes under both categories are structure molecule activities **(A-1)**, catalytic activities **(A-2)**, and binding activity **(A-3)**, cellular metabolism **(B-1)**, ubiquitin-dependent proteolysis activities **(B-2)**, and cell organization and biosynthesis activities **(B-3)**.

### Most of QTL-Ub genes are regulated during PCW/SCW transition stage

Both expression and functional analyses indicated ubiquitin-pathway associated genes as major players in the development of Pima fiber. Applying reiterative BLAST searches using consensus motifs for plant Ub/26S-proteasome genes identified ~ 650 genes encoding Ub/26S proteasome-related genes in the fiber transcriptome. This represented 7.60% of the fiber transcriptome. Out of these 650 genes, only 241 were homologous to ubiquitin-related gens in *Arabidopsis*. Approximately 74.92% (487 genes) and 17.38% (113 genes) of ubiquitin-and 26S Proteasom-related genes, respectively, were printed on the microarray platform used in this study. Only 25.66% were developmentally regulated in the fiber transcriptome (Table 2). However, 50% of the QTL-associated Ub-pathway genes (QTL-Ub) were developmentally regulated (Table 2). This ratio is 2-times higher than the expected global ratio of significance, which is estimated at 23.99% in this study. The majority of the developmentally regulated QTL-Ub genes were members in the cellulose synthesis and PCW/SCW transition drQTLs. As illustrated in Figure 6, the dynamics of the 15 QTL-Ub genes undergo a significant change in their expression pattern during PCW/SCW transition stage. E3 ligases were the most represented family among the three categories of the Ub-pathway genes (Table S4). In the QTL-E3 ligases, genes with RING finger domains were highly represented (Table S4). Three developmentally regulated ubiquitin-related genes representing the three families involved in this pathway (E1, E2, and E3) were associated with three fiber fineness QTLs on chromosomes 6, 15, and 17 (Table S4), and were co-expressed in a similar pattern, which reflected an abrupt increase in the expression level at the transition stage PCW/SCW relative to both PCW and SCW (Figure 6). AtCUL1 (E3 ligase) that is linked to FF15_1 QTL was highly up-regulated with a steady increase in the expression level from PCW through SCW stages. Given the dependence of fiber fineness on the duration of the PCW/SCW transition stage, this finding strongly suggests that UBQ1 [FF15_1], RING finger [FF6_1], and E3-dpendent [FF17_1] genes are key factors in the development of fiber fineness superior trait via regulating the transition stage PCW/SCW. A RING finger gene mapped to FS_25 was up-regulated only during the PCW/SCW transition stage. The highest expressed QTL-Ub gene was an F-box E3 ligase, which was mapped to fiber elongation (FE F-box) QTL at chromosome 23. Another two E3 ligases, SKP1-like and AtCUl1, were mapped to FF QTL on chromosomes 5 and 15, respectively, and were expressed in a similar pattern to FE F-box gene. These three genes were expressed at same level during PCW followed by a big increase in the expression during both PCW/SCW and SCW stages, which suggests strong correlation with the transition and cellulose deposition stages in the developing Pima fiber.

**Figure 6:**
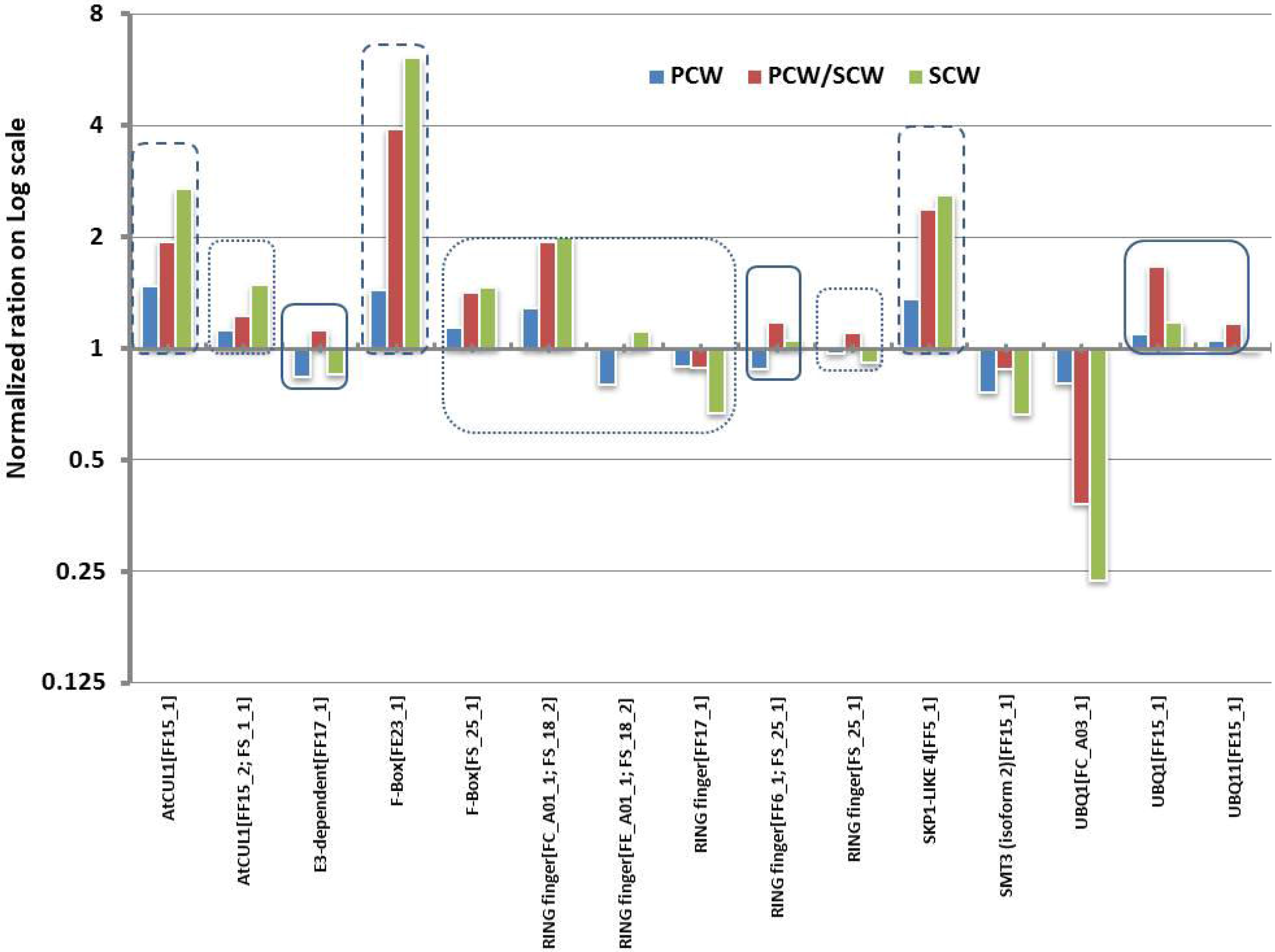
Average relative expression ration of QTL Ubiquitin-related genes during PCW and SCW development of Pima fiber. The ratio SCW/PCW shows that there is a significant increase in the expression level in association with the transition from PCW to SCW developmental stages. Notice that Contig ID is abbreviated however the complete ID is mentioned in Tables S5 and S6.

### Interaction between UBC9 and elongation related genes

Yeast-two-hybrid screens (Y2H) using UBC9 as bait and the transcriptome of developing Pima fibers at 14-17dpa revealed several interesting interactions (Table 3). Reciprocal interactions were identified by using prey genes as baits. Among the identified prey genes are E3 ligase CHIP, Dynamin, Arginase, and β-tubulin.

## Discussion

Expression profiling of the transcriptome and quality QTLs of Pima fibers during the morphogenesis revealed genes and drQTL networks directly linked to the development of different fiber traits. These results provided deeper insights into the genetic programs controlling the development at each stage and contributed to building a model for Pima fiber developmental.

### Pima fiber development model: expression to phenotype

The proposed model (Figure 7) integrates three levels of information in the fiber development to provide a deeper understanding of genetic programs underpinning the development of Pima fibers: developmental time frame, developmental expression profiles, and drQTLs networking. It is important to keep in mind that drQTLs were identified as an association between developmental gene expression and fiber QTLs.

**Figure 7:**
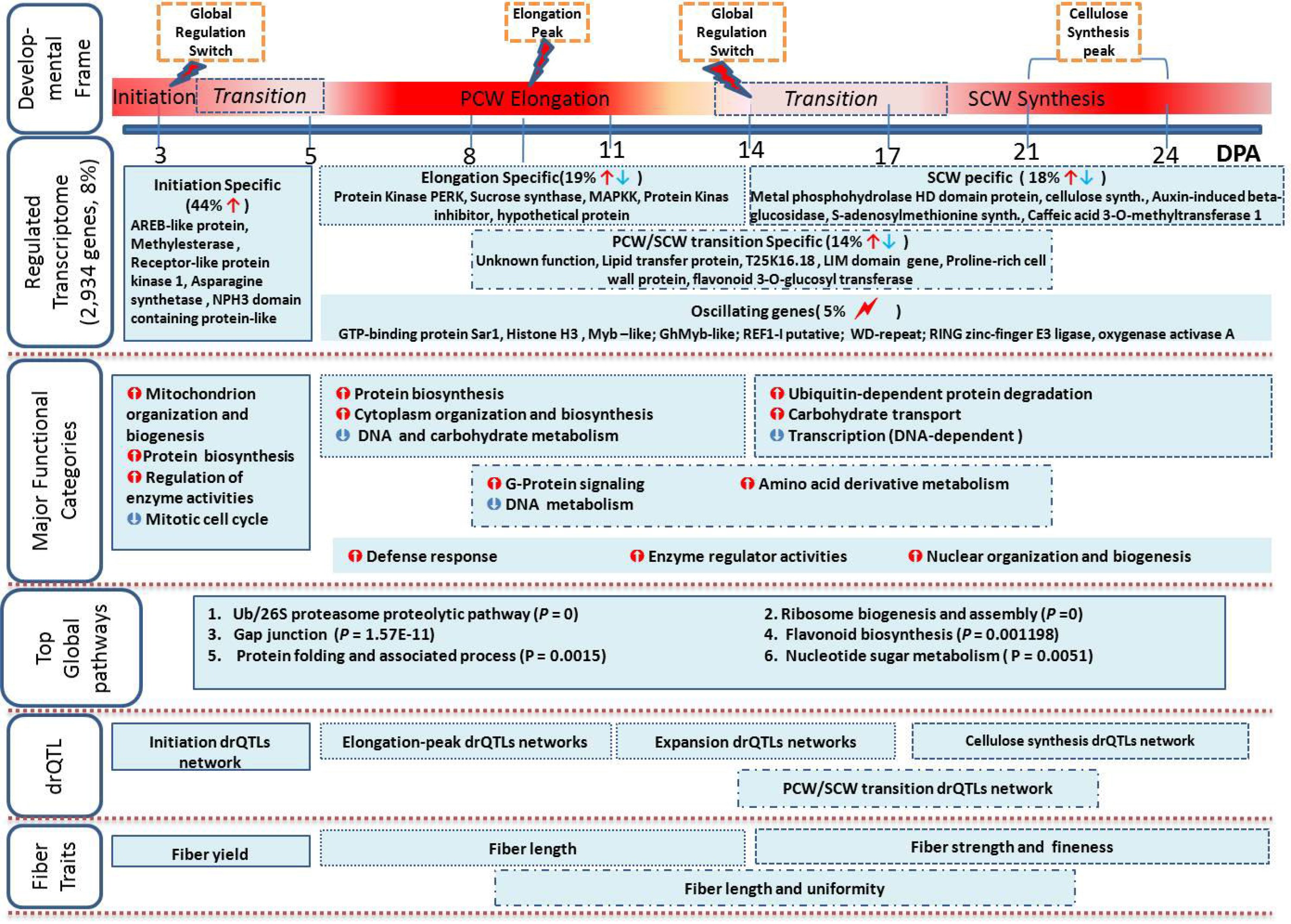
A schematic model for the transcriptional and genetic activities that underpin the development of Pima cotton fiber.

### Developmental time frame

According to the classical model of cotton fiber morphogenesis [8], fiber development occurs though 4 overlapped stages. The timing and duration of each fiber developmental stage depends on both genotype and environmental conditions. Pima fiber growth rate enters a linear phase of cellular elongation at 5dpa and reaches expansion peak around 9.5dpa (2.3 mm/day). Fiber attains almost its final length by 21dpa (Figure 1). Therefore, these 3 points are cornerstones in the developmental frame where they represent the beginning of PCW accentuated growth, peak of cellular elongation, and end of the elongation stage accompanying by commencing of SCW deposition, respectively. The transition between PCW and SCW (PCW/SCW) is usually an overlooked stage. Extra-long staple genotypes have a greater overlap between fiber elongation and SCW biogenesis stages than other genotypes do [13, 43–45]. In this study, there are 3 lines of evidence that PCW/SCW stage starts at 14dpa and ends around 21-24 dpa: the steady decrease in fiber length from 14 to 21dpa (Figure 1F), the global switch in the expression at 14dpa of all developmental profiles (Figure 2), and the abruptly increased number of significantly regulated genes at 14 through 21dpa. During the PCW/SCW stage, expansion, cellulose depositions, expansion cessation, and cytoskeleton re-orientation are the major processes. These steps are important in defining the quality of fiber traits. Under the same growth condition, Pima fiber growth rate is faster than upland and it reaches peak one day before upland [28]. The development of Pima fiber does not only require longer time but also undergoes faster growth program.

### Developmental expression profiles

In the proposed model (Figure 7), 44% of the regulated transcriptome was down-regulated after 5dpa, indicating that this portion contributes to the early-expansion or transition between initiation and elongation (initiation/PCW: ~3-5dpa) stages and possibly to the initiation (~-3 to 3dpa) stage. This percentage refers to the complexity and diversity of the biological processes taking place during these developmental stages, which in turn control fiber yield. At the transcription regulation level, AREB transcription factor might be a key player during the initiation and initiation/PCW stages. AREB-like protein is an ABA responsive element binding protein. Transgenic plants overexpressing the phosphorylated active form of AREB1 expressed many ABA-inducible genes such as RD29B, without ABA treatment, which suggested that ABA-dependent phosphorylation of AREB1 regulates its own activation in plants [46]. Given that ABA accumulation inhibits fiber elongation [47, 48], the confinement of AREB-like protein to initiation/PCW stage might play a major role in controlling the beginning of elongation stage in a way that allows enough time for initiation and synchronize the starting of elongation stage. This hypothesis can be supported by knowing that the down-regulation of AREB1 at 5dpa (Type-1 genes, Figure 2) is associated with the beginning of elongation stage. During the initiation and initiation/PCW stages, mitochondrion organization and biogenesis, and protein biosynthesis activities were top represented functional categories. Mitotic cell cycle was the most under represented activities during the initiation/PCW stage, which is expected because fiber cell does not undergo proliferation. Mitochondria play a central role in energy metabolism and therefore affect the survival and proliferation of the cell. Since fiber cell does not proliferate, the increased activities of mitochondrial organization are possibly related to the transition from initiation to PCW stage. Studies have shown that mitochondria are morphologically dynamic in such a way that their shape and distribution in the cell can affect the response to different stimuli [49], which in turn will have an influence on the duration and speed of the developmental stages.

During the elongation stage, 19% of the developmentally regulated genes control the elongation stage (Type-2 genes and Figure 2). Fiber elongation is the process of unidirectional increase in length due to the coordination between two major biological processes: turgor pressure and primary cell wall loosening and expansion. Conceptually, several metabolic processes are involved in building and elongating the plant primary cell wall: synthesis of new wall materials, secretion, transportation, integration and assembly, and stress relaxation and polymer creep [50]. In accordance with this, protein biosynthesis and cytoplasm organization and biosynthesis are the most over represented processes during the elongation stage as deduced from this work (Figure 7). Gene families such as tubulin, expansin, and Susy are classically known to be elongation-associated. However, the active members/isoforms are different from cell to cell and between genotypes. In our model for Pima fiber development, β-tubulin isotypes are more abundant than a-tubulin isotypes during the polar elongation. Also, α-expansins are the major loosening enzyme during same stage. New candidate gene families active in the elongation process of Pima fiber were identified. For instance, a proline-rich extension-like receptor protein kinase (PERK, more than 8 folds), Mitogen-activated protein Kinase (MAPK, more than 4 folds) were in the top 5 up-regulated genes during the elongation stage of Pima fiber. These 2 genes are member of the Kinase family, which plays key regulatory roles in all aspects of eukaryotic cell physiology [51, 52]. Beginning after 14dpa in the proposed model (Figure 7), 18% of the developmentally regulated transcriptome contributed to the biogenesis of SCW stage, which is dominated by cellulose synthesis and deposition [12]. This stage has major influences on fiber strength and elongation that are controlled by the thickness of cellulosic chains, fibrillar and crystalline structure, and the extensive inter-and intra-molecular hydrogen bonding [53], while fiber fineness is a function of both length and thickness. In *Arabidopsis*, cellulose biosynthesis is catalyzed by the rosette of cellulose synthases, which is composed of six subunits, and each subunit is composed of six cellulose synthases [54]. In the cotton fiber transcriptome there are eight different members of *CesA* and *CesA*-like (*Csl*) not including *GhCesA1* and *GhCesA2* that have been characterized from *G. hirsutum* [55]. Surprisingly, all *CesA* genes except one were expressed at low levels during development, which suggests they are all associated more with the biosynthesis of PCW cellulose. The only up-regulated cellulose synthase subunit during SCW (15.65 fold) was highly similar to the *Gossypium hirsutum* cellulose synthase-like catalytic subunit (*CelA3*) (Table S2). In association with cellulose synthase, endo-1,4-beta-glucanase (EGase family) is also up-regulated during SCW biosynthesis (Table S2). EGase is possibly involved in trimming off defective glucans from growing microfibrils or cleaving off the sitosterol-beta-glucose primer from the growing glucan chains [39]. In the fiber transcriptome, there are about 10 members of the EGase family. Two major isoforms were co-expressed (19.97 fold) with *CesA* during the synthesis of Pima fiber SCW. One more EGase (Korragen) gene was about 4 fold up regulated during the cellulose synthesis stage. Ubiquitin-dependent protein degradation and carbohydrate transport functional are the most over-represented categories during SCW biosynthesis in the proposed model (Figure 7). Regulated protein degradation through Ubiquitin conjugation plays a crucial role in all aspects of development [56]. This pathway will be discussed separately. It was not unexpected to find carbohydrate transport activities over-represented during SCW synthesis given the domination of cellulose synthesis, which creates more demands on carbohydrate (sugars) substrates and the subsequent transport to the plasma membrane where cellulose synthesis takes place. However, in both plants and microorganisms, sugars levels in the cell affect the sugar-sensing systems that initiate changes in gene expression [57]. Therefore, the over-representation of carbohydrate activities can be viewed as a gene regulation mechanism in the developing Pima fiber. Although it is well established that cellulose synthesis and deposition are the dominant activities during SCW biosynthesis in developing fiber, there clearly is a diversity of genes/processes and metabolic activities taking place, associated with cellulose biosynthesis.

### Potential drQTL networks

QTLs are genomic regions associated with phenotypic variations of a trait within a population; nevertheless these regions may not harbor genes controlling the associated phenotypic variations [58]. Expression QTLs (eQTLs) are an approach to explore the relationship of phenotypic variations with transcriptome variations by employing microarrays to profile the global gene expression across individuals from segregating population followed by mapping the variance of association [19, 59]. Although informative this approach is very expensive, requires extensive experimental work, and does not use the available association data that have been generated, and still ongoing, over the last two decades. Herein an indirect approach to identify transcription variations associated with phenotypic variations is described. In this approach, developmental transcriptional profiling of previously mapped QTL-genes for different fiber quality traits was performed and followed by clustering to identify co-expressed QTL-genes, which are labeled as developmentally regulated QTLs (drQTLs). The lack of fully saturated QTL maps is a drawback of this approach; however the unstopped efforts in sequencing full genomes will overcome this problem. Five drQTLs networks are identified; each of them is corresponds to a major stage in Pima fiber development. Interestingly, gene members of each drQTLs network are physically mapped to QTLs for different traits. For instance, on the linkage group A01 and its homoeologous chromosome (Chr. 18), a group of 15 early-expansion regulated genes were mapped to 6 overlapped QTLs: FC_A01_1, FF_A01_1, FF_A01_2, FL_A01_1, FS_A01_1, FS18_1, and FS18_2 (Table 3). The same idea is true with all five drQTL, which suggests that QTL associated genes interact and play important roles in fiber development at various developmental stages influencing various fiber traits, and are not limited to genes mapped to particular QTL that is associated with specific trait. Each drQTLs network has unique and overlapped members. The features and characteristics of the identified drQTLs are of great accordance with results from regular eQTL approach [58, 60, 61]. Although the saturation level of the consensus QTL is relatively low, key genes and pathways were identified in each of the drQTLs. For instance, cellulose synthesis and PCW/SCW drQTLs networks were found enriched with Ubiquitin pathway associated genes, which led to hypothesizing a significant role for this pathway in regulating the transition stage PCW/SCW.

### The Ub-pathway is a potentially regulating the PCW/SCW transition stage

Regulated protein degradation by ubiquitin/26S proteasome contributes significantly to development by affecting a wide range of processes, including embryogenesis, hormone signaling, and senescence [56]. Collectively, the Ub-pathway accounts for 7.4% of the fiber transcriptome comparing to 5% in the whole *Arabidopsis* transcriptome, and it is the most over-represented pathway in the developmentally regulated Pima transcriptome. Specifically, developmentally regulated Ub-pathway genes are preferentially linked to fiber quality QTLs and the majority of Ub-QTL genes are differentially regulated during the transition stage PCW/SCW (Table 2). Moreover, cellulose synthesis drQTLs and PCW/SCW transition drQTLs are enriched with Ub-pathway genes. Finally, the majority of Ub-QTL gene members in the PCW/SCW transition drQTL network are mapped to fiber fineness QTLs; this is an exciting finding knowing that fiber fineness trait is a function of both fiber length (PCW) and thickness (SCW), and hence is influenced largely by the transition stage. To identify the mechanism of regulating the transition PCW/SCW, it would be crucial to identify what specific proteins are targeted for degradation using Ub-QTL genes. Y2H screens using UBC9 as bait and Pima total RNA from the PCW/SCW transition stage (14-17dpa), Identified several interactions, including E3 ligase CHIP, Dynamin, Arginase, and 3-tubulin. These genes were shown to be associated with cellular elongation, polar expansion, plasma membrane remodeling, cytoskeleton dynamics and basic amino acids transfer [62–65].

In conclusion, this study provides novel insights into the genetic factors underpinning the development of Pima fibers as well as reinforces of previously characterized fiber related genes. In the light of identified developmental time frame, the identified developmental gene Types (Type-1 to 5) and the proposed fiber quality drQTLs provides an advanced stage in explaining the genetic program of each stage in Pima fiber development and hence the dependent phenotypic traits.

## Materials and Methods

### Plant Material

Cotton (*Gossypium barbadense* L. cv. Pima S7) was grown under optimal conditions in the greenhouse under a 30/21°C day/night temperature regime in a randomized complete block design to generate two biological groups (g-A and g-B). Flowers were tagged on the day of anthesis (0 dpa), and developing bolls from the first fruiting position closest to the stem were collected at 5, 8, 11, 14, 17, 21, and 24 dpa between 8 am and 11 am to eliminate diurnal effects. Cotton bolls (5-6) from the same developmental stage were collected from individual plants, and pooled together in 50 ml polypropylene tubes filled with liquid nitrogen and stored at −80°C. Immature fibers were isolated from more than one pool in liquid nitrogen as described [10] and stored at −80°C. A growth curve was drawn for developing cotton fibers by measuring fiber length (mm) from a minimum of 10 locules representing 10 bolls from 10 different individual plants as follows. Locules were boiled gently in ddH2O for ~15-20 min to untangle fibers and stained in an aqueous solution of 0.05% methylene blue for 1-2 min. Fibers from individual ovules were gently spread on a watch glass, and three independent fiber length measurements were determined per ovule, measuring from the ovule surface to the fiber tip using a digital caliper. Fiber length measurements were made from 30 ovules (3 ovules/locule) per developmental stage to generate 90 data points for each developmental stage. Fiber growth rate (mm/day), and the Standard Error Rate (ER) at seven developmental time points were calculated.

### Microarray hybridization

Total RNA was isolated independently from each biological sample as technical replicates using the hot borate method [36, 66] following modifications described by Arpat *et al.*, (2004) [6]. The quantity and integrity of the RNA was assessed spectrophotometrically and by agarose gel electrophoresis in 1% agarose/6% formaldehyde/1X MOPS gels stained with SYBR gold nucleic acid gel stain (Molecular Probes) in 1X MOPS buffer following electrophoresis [67]. Total RNA (10 μg) spiked with 1000 pg of a reference mRNA spike mix (Lucidea Universal Scorecard, Amersham Pharmacia) was used to synthesize first-strand cDNA in the presence of amino allyl-modified dUTP (aa-dUTP) [6, 68]. The reverse transcription reaction mix contained ~10 μg “spiked” total RNA, 5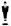g poly-T21VQ primers (Operon), 0.2 mM dNTPs (Sigma), 0.2 mM aminoallyl-dUTP/dUTP (Sigma) in a 2:3 ratio, and 2.5 U SuperScript III (Invitrogen). Following incubation for 2 hours at 45°C, the aminoallyl-labeled cDNA was purified using Microcon YM-30 columns (MilliPore) according to manufacturer’s instructions. Aminoallyl-cDNA was labeled using Alexa Fluor 555 and 647 succinimidyl ester dyes according to manufacturer’s instructions (Molecular Probes, Inc.), and purified using the QiaQuick PCR purification kit (Qiagen).The used cotton oligonucleotide microarrays platform (NCBI-GEO: GPL6917) carried 12,227 unigenes from the cotton fiber EST database, 71 negative and positive control elements, transgene and vector controls, 6 spike-in controls (Alien Spike mRNA, Stratagene), and blank and buffer controls were spotted on the microarray slides (Corning GAPSII slide) in duplicate and arranged in 23 X 23 subarrays [6]. Printed slides were pre-hybridized in 5X SSC, 0.1% SDS, and 1 mg/μl BSA in a sealed coplin jar for 45 minutes at 42°C, followed by two washes of 10 sec each in filtered ddH_2_O preheated to 42°C, and dried with filtered compressed air. Alexa-labeled cDNA (500 ng/fluor or the equivalent of 40 to 45 pmol of dye depending upon the frequency of incorporation (FOI)) was suspended in 45μl of hybridization buffer (25% formamide, 4.16X SSC, 0.08% SDS, and 0.1 μg/pl sheared salmon sperm DNA) and loaded underneath the cover-slip (M Series LifterSlip, 22x60mm, Erie Scientific) using capillary action. Following hybridization at 42°C for 16-18 hours in a humidified hybridization chamber (Genetix), slides were washed sequentially at 42°C in 2X SSC, 0.2% SDS for 5 min, 0.5X SSC for 3 minutes, 4 times in 0.1X SSC for 1 minute per wash, and 0.05X SSC for 10 seconds, followed by drying with filtered compressed air.

### Microarray Experimental Design and Data Analysis

A bi-directional double loop hybridization [69] with dye swap experimental design encompassing a total of 28 hybridizations was employed to maximize discovery of significant changes between developmental stages. Self-hybridization controls were conducted to evaluate both within-array correlation and dye bias [28]. Hybridized slides were scanned using a GenPix3000 array scanner (Axon Instruments, CA, USA) followed by feature extraction to quantify signal intensities using ImaGene 4.2 software (BioDiscovery). Data were filtered and normalized, quality assessed, and statistically analyzed as described in detail in Alabady *et al.*, (2008) [28].

### Cluster, function, and pathways analysis

The expression profiles were developed by clustering significantly expressed genes according to the expression similarities using K-means clustering GeneSpring (Agilent). Parameters selected for clustering were K = 10, standard correlation as a similarity measure, and 100 iterations. A K = 10 was chosen after testing a series of K values (5, 7, 10, 14, and 16), which accounts for more than 90% of the variability within the data with minimum number of clusters. Functional analysis of significantly expressed genes was performed using GenMapp software (www.genmapp.org). The over-/under-represented GO annotation were identified within each cluster of similarly expressed genes using GO annotation (www.geneontology.org) followed by MAPPFinder function in GenMapp software [70]. Redundant GO results with *p* <0.01 were normalized to select for the outermost possible parent GO term. Pathways analysis was performed using KGG Orthology-Based Annotation system, KOBAS [71, 72].

### Expression analysis of fiber QTLs

A bioinformatics approach was used to associate significantly expressed candidate genes with fiber QTL data. Fiber QTLs data were taken from the primary QTL map [18] for the following reasons: (1) a large number of fiber QTLs mapped to one population with a strong LOD, (2) QTL associated markers were mapped to the linkage most saturated cotton linkage map [3], and (3) one of the parent of the mapping population was Pima, which is the main focus of this study. A Consensus QTL map was developed by employing a custom PERL script that links QTL loci from the primary map to corresponding loci on chromosomes and homoeologous chromosomes in the genetic linkage map [3]. The conjoined STS/QTL gene lists were cross-linked to the differentially regulated genes in the developing Pima fiber transcriptome by BLAST. Differentially regulated QTL associated genes were clustered using K-means to identify stage specific QTL expression profiles.

### Expression analysis of Ub-associated QTL genes

Ubiquitin-dependent proteolysis pathway genes in the fiber transcriptome were identified by blasting cotton fiber unigenes against Ubiquitin pathway (Ub) database (http://plantsubq.genomics.purdue.edu/). Fiber Ub-genes were classified further based on whether they were significantly expressed and temporally regulated, and if the candidate genes were also associated to any fiber QTL for color (FC), elongation (FE), fineness (FF), length (FL), length uniformity (FLU), and strength (FS). The expression profiles of the differentially regulated Ub-QTL genes were linked to the fiber developmental stages.

### Validation of microarray results

Relative gene expression was quantified for 25 selected genes (Table S6) using real-time PCR (q-RT-PCR) to validate microarray results by normalizing expression of each gene to a non-regulated reference gene [73]. A constitutively expressed actin gene that showed stable expression across all developmental stages was selected as an internal reference for this study. Gene-specific primers (GSP) were designed using custom Perl scripts, which uses the Primer3 algorithm and applies high stringency specification and eliminates cross-hybridizing primers by blasting against all known cotton sequences (approximately 52,000 genes) (Tables S5 and S6). Total RNA (5 ug) was treated with RNase-Free DNase (Promega) to remove trace levels of contaminating genomic DNA. DNA-free RNA samples were reverse transcribed using SuperScript III in a mix of both oligo(dT) and random hexamer primers (1:1). Real-time PCR reactions were performed in triplicate using the SYBR-Green PCR Master Mix (Applied Biosystems) on an ABI 7000 instrument (Applied Biosystems). An assumption-free analysis of q-RT-PCR data [74] was applied to estimate PCR amplification efficiencies for each gene. The efficiency was calculated using LinRegPCR software [74]. Data analysis was performed with a relative quantification software tool, REST, [75] using the geometrical mean of actin expression as an internal reference. This software corrects the results for target and reference genes according to their calculated PCR efficiencies, and compares the levels of transcript present in different samples by performing a pair-wise fixed reallocation randomization test to determine the statistical significance of the changes in transcript abundance.

## Ubiquitin interactions using Y2Hybrid experiment

Y2H experiment was conducted suing Clontch Matchmaker Gold Yeast Two-Hybrid System following the manufacturer recommended procedures.

## Data release

Raw and processed data obtained from this study were deposited in MIAME compliant format in NCBI-GEO (GSE 11689).

## Acknowledgment

The authors would like to thank Professor Thea A. Wilkins, who is now retired, for enabling us to conduct this research in her laboratories at University of California-Davis and Texas Tech University.

## Supplementary Material

**Figure S1:**
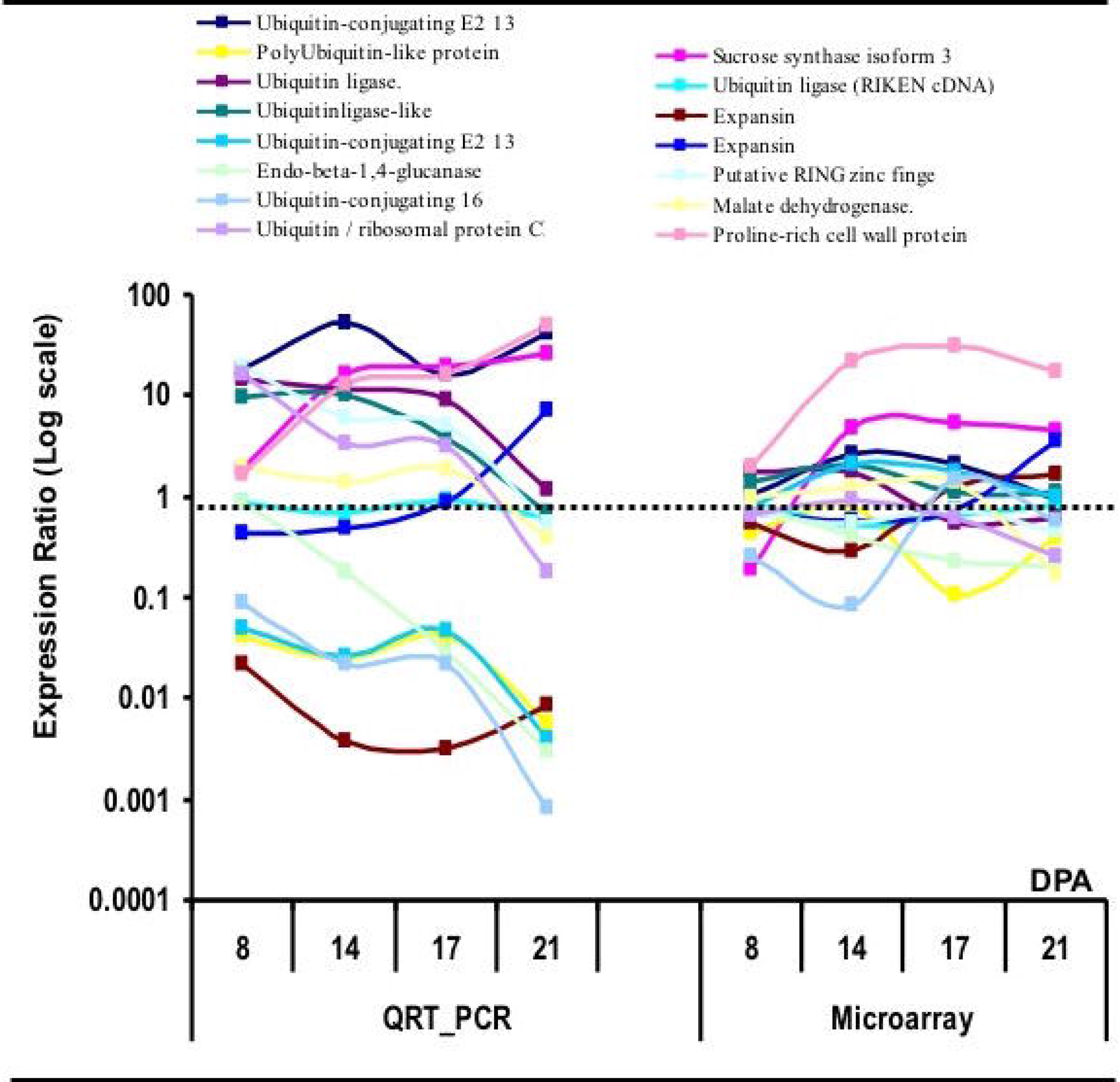
Validation of microarray data. The expression of randomly selected genes were quantified at 4 time points (dpa) using SYBR-green QR-PCR. The QRT-PCR-based and microarray-based expression profiles showed similar patterns of expression dynamics in most of the cases (12 genes out of 15).

**Table S1**: List of all significantly expressed genes (and expression data)

**Table S2**: Top regulated genes during the polar elongation and SCW synthesis of growing Pima fiber cell. Expression during both polar elongation and SCW was calculated the average of log2 ratio at 8, 11 and 14 dpa, and at 17, 21 and 24 dpa, respectively. The cluster number refers to which of the 10 k-means cluster each gene was assigned.

**Table S3**: Percentages of significantly expressed genes of whole microarray set, QTL associated genes and QTL-Ubiquitin genes during Pima fiber morphogenesis.

**Table S4**: GenMapp annotations and pathways profiler results showing over (+ Z score) and under (− Z score) represented annotations and pathways within the 10 clusters of fiber transcriptome.

**Table S5**: List of the QTL-ubiquitin genes that were differentially expressed genes during the development of Pima fiber.

**Table S6**: Genes and primers used to validate microarray results using quantitative real time PCR (qRT-PCR)

